# Modeling suggests that multiple immunizations or infections will reveal the benefits of updating SARS-CoV-2 vaccines

**DOI:** 10.1101/2022.05.21.492928

**Authors:** Rajat Desikan, Susanne L. Linderman, Carl Davis, Veronika Zarnitsyna, Hasan Ahmed, Rustom Antia

**Author notes:** Correspondence (R.D.), (R.A.). These authors contributed equally (Rajat Desikan, Rustom Antia).

## Abstract

When should vaccines to evolving pathogens such as SARS-CoV-2 be updated? Our computational models address this focusing on updating SARS-CoV-2 vaccines to the currently circulating Omicron variant. Current studies typically compare the antibody titers to the new variant following a single dose of the original-vaccine versus the updated-vaccine in previously immunized individuals. These studies find that the updated-vaccine does not induce higher titers to the vaccine-variant compared with the original-vaccine, suggesting that updating may not be needed. Our models recapitulate this observation but suggest that vaccination with the updated-vaccine generates qualitatively different humoral immunity, a small fraction of which is specific for unique epitopes to the new variant. Our simulations suggest that these new variant-specific responses could dominate following subsequent vaccination or infection with either the currently circulating or future variants. We suggest a two-dose strategy for determining if the vaccine needs updating and for vaccinating high-risk individuals.

## Introduction

SARS-CoV-2 (‘CoV-2’ hereafter) has caused the most severe pandemic since influenza in 1918 (approximately half a billion confirmed cases and 6 million deaths as of 28^th^ April 2022 - WHO). In contrast with the 1918 influenza pandemic, where no vaccines or therapeutics were available and immunity was only gained following recovery from infection, vaccination has played a key role in mitigating the morbidity and mortality of CoV-2 (*1, 2*). However, as is the case with other circulating human coronaviruses, immunity does not provide lifelong protection from reinfection (*3*–*5*) and we are witnessing waves of infection with new virus variants. These variants arise and spread due to a combination of factors such as waning immunity (*6*–*9*) and virus evolution (*10*–*12*). The latter results in both more transmissible viruses (*13*–*16*), and viruses able to escape immunity to earlier variants and vaccines (*16*–*18*). In particular, the Omicron (OM) variant of CoV-2, that arose in late 2021, is much more transmissible than the ancestral Wuhan (WU) (*13, 15*), and in addition, OM has a panoply of mutations in the spike protein (*12, 16*) that allow it to partially escape antibody responses to earlier variants as well as Wuhan (WU) based vaccines (*2*).

Prima facie, we might expect that it is best to keep the vaccine updated with the current strain. For example, we might expect the updated vaccine to generate higher antibody-titers to the currently circulating virus in unvaccinated individuals. Indeed, the experimental data from the animal model studies support this (*19*). However, over time, most of the population will have either been immunized or naturally infected. Studies on influenza have shown that prior immunity can skew responses to subsequent infection and immunization and the phenomenon has been termed original antigenic sin (OAS) (*20*–*25*). Understanding of the implications of OAS for CoV-2 vaccination requires integrating experimental and clinical studies with mathematical models. A number of elegant experimental and observational studies show that prior immunity has unexpected effects on the outcome following boosting with different vaccines (*26*–*28*), and in particular suggest that updating the vaccine to match the circulating variant does not enhance the antibody titer to the circulating variant any more than the original vaccine. We focus on the OM-vaccine study by Gagne et al. (*28*) as the pattern of boosting observed was very similar to the study using the vaccine based on the Beta variant (*27*).

The Gagne study (*28*) used a macaque primate model system to compare the boosting of immunity with a WU-versus an updated OM-vaccine. Primates were first given two vaccine doses of the currently used mRNA-1273 vaccine (WU-vaccine), which encodes a spike protein derived from the ancestral Wuhan virus variant, to mimic prior immunity of vaccinated humans. These two vaccinations (#1 and #2) resulted in a high titer of antibodies against the WU virus, and significantly lower titers against the OM variant (see Fig 3A). Over time the antibody titers to both WU and OM viruses waned significantly, and at week 41 the animals were boosted with a third vaccination, either with the original WU-vaccine (vaccination regime WU-WU-WU) or an updated OM-vaccine that incorporated spike protein antigen from the OM virus (regime WU-WU-OM). This allowed them to determine whether updating the vaccine would produce higher titers to the OM-virus. Surprisingly, their results showed that both WU-WU-WU and WU-WU-OM resulted in similar antibody titers to the OM-virus. Also surprising was the finding that both these vaccination regimes resulted in similar antibody titers against the WU-virus (albeit at higher levels than to the OM-virus, as shown in Fig 3A). These observations implied that it might not be necessary to reformulate the vaccine to match the OM virus variant.

In this paper, we use computational models to better understand the rules of boosting of responses to new virus variants. We develop and use computational models to analyze the results of Gagne et al. (*28*) as well as data from other CoV-2 (*8, 19, 29, 30*) and influenza (*31*) studies. We show that our model can capture the key features of the boosting of immunity following immunization with the WU- and OM-vaccines. Analyzing the dynamics of antigen, B cells, and antibodies in our simulations allows us to understand the reason for the initially surprising observation that vaccination #3 with either vaccine results in similar antibody titers to the OM virus. We then use this model to explore what might happen following subsequent vaccinations. We find that while the level of immunity to the WU and OM viruses appears equal following the initial booster with either the WU- or OM-vaccines, using the OM-vaccine may have significant advantages with subsequent vaccinations or infections. Based on model predictions, we suggest critical experiments that will allow us to determine whether the vaccine strain should be updated to that of the circulating virus variants.

## Results

### The immunodynamics model

We consider an immunodynamics model for the interaction between the vaccine and the humoral immune response. The model extends earlier multi-epitope models for the dynamics of antibody levels following vaccination (*25, 32*) in the following ways. First, we incorporate two different vaccines, the WU- and OM-vaccines. Second, we incorporate differences in the boosting of naïve and memory cells to antigenically altered epitopes that underlie the phenomenon of original antigenic sin (*23*). We then used the model to explore how the boosting of immune responses to the new virus variants is affected by the interplay between prior-immunity to the old variant and the antigens expressed by the updated vaccines.

The model is shown schematically in Fig 1. The WU- and OM-vaccines have unique as well as shared or cross-reactive epitopes. We keep track of three types of epitopes: C, W and O denote cross-reactive epitopes and epitopes unique to WU- and OM-vaccines, respectively. We also keep track of B cells and antibodies specific to these epitopes. B cells specific to an epitope are stimulated by cognate antigen, undergo clonal expansion, and produce antibodies specific to that epitope. The response wanes once the antigen is cleared. Further details, equations and parameters are described in the Materials and Methods. We do not include more complex features of the selection and differentiation of B cell clones and interactions with other immune cells such as follicular dendritic cells and T cells in germinal centers (*33, 34*). This is because, at this stage, the experimental data does not include precise measurements of these quantities after CoV-2 vaccination. Under these circumstances, the results of simpler models can typically be more robust than those of complex models (*35*), and we focus on qualitative patterns observed in the data rather than specific values.

**Fig 1:**
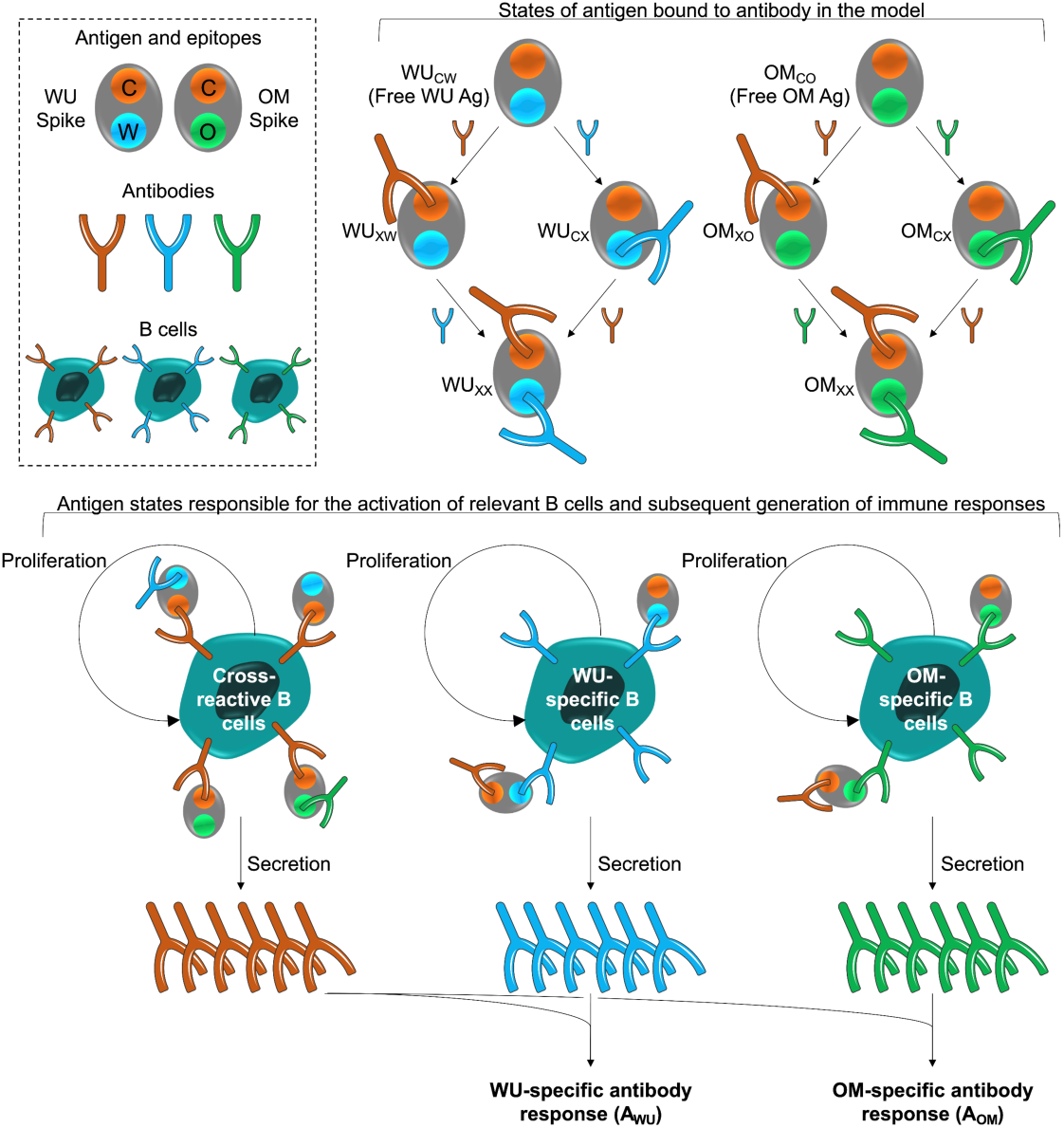
Model schematic. The box at the top left shows the epitopes of the WU- and OM-vaccines. Epitope C (shown in orange) is common to both vaccines. Epitopes W (blue) and V (green) are unique to the WU and OM respectively. Antibodies specific to these epitopes can bind to these epitopes and prevent them from stimulating B cells for the same epitope. The different antigen states generated and the B cells they stimulate are shown in the top right and bottom panels respectively. The bottom panel illustrates that binding of antigen to B cells stimulates their clonal expansion and the production of antibodies.

### Model recapitulates a number of studies on CoV-2 responses following vaccination and boosting

Our model recapitulates the broad patterns of immunity generated both by natural infections and vaccination with CoV-2. A wealth of data show that both natural infection with circulating CoV-2 as well as vaccination induce antigen-specific humoral immune responses. We next describe how the model can qualitatively describe the pattern of the humoral immune response observed in a number of studies.

As mentioned in the Introduction, prima facie we would expect that boosting of naïve individuals with a vaccine based on the circulating variant will elicit higher antibody titers to this strain rather than a vaccine based on an earlier variant. This simple observation was demonstrated by Ying et al. (*19*) as seen in the left panel of Fig 2A. In their experiment, groups of mice were immunized with two doses of either the WU-vaccine (WU-WU) or the OM-vaccine (OM-OM), and the generated WU and OM antibody titers were compared between the groups. The WU-WU group elicited orders of magnitude higher WU titers than OM titers, while the OM-OM group exhibited exactly the opposite response, much higher OM titers than WU titers. Our model recapitulates this observation.

**Fig 2:**
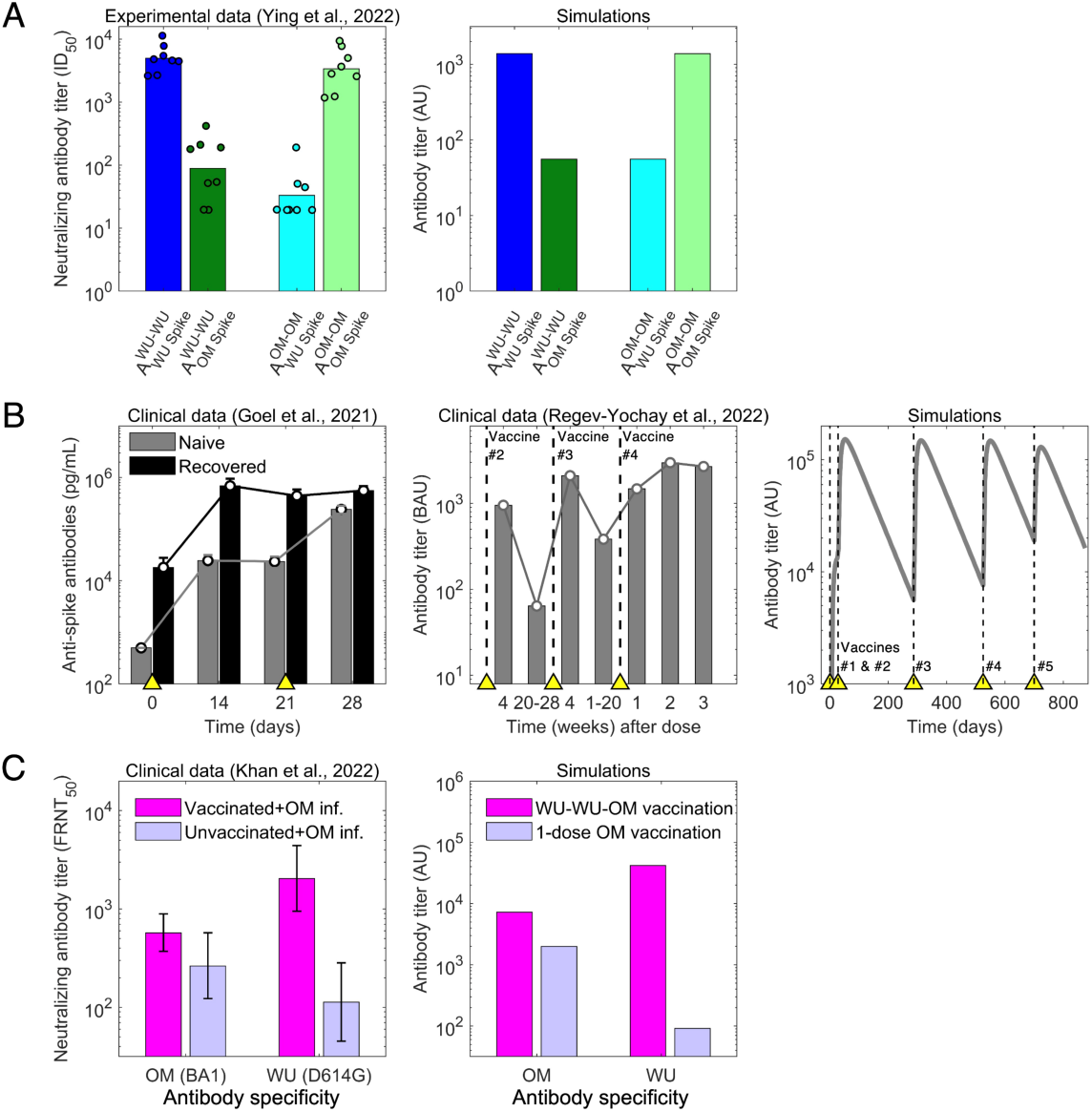
The model recapitulates antibody responses to CoV-2 following vaccination and infection. **(A)** We show data for immunization of mice with 2 doses of either the WU- or the OM-vaccine. We see that high antibody titers to a given antigen (WU or OM) requires 2 vaccinations incorporating that antigen. **(B)** Our model recapitulates the saturation of antibody titer following repeated immunization or infection. The left panel shows data for the virus titer in naive (grey) and CoV-2 infected and recovered (black) individuals following two doses of the WU-vaccine. We see that the titer of antibodies in recovered individuals saturates after a single vaccination, while that in naive individuals is boosted by the second vaccination. The middle panel shows data following four doses of the WU-vaccine and shows that the virus titer is boosted and reaches a plateau after vaccination #3. Simulations of repeated immunizations with the WU-vaccine at times indicated by the yellow triangles show that the antibody response to the WU-virus increases substantially after the first two vaccinations. Further boosts with the same vaccine results in little further increases in the titer of antibody. **(C)** We plot data from a human study showing that OM infection causes higher antibody titers to OM compared with WU in unvaccinated individuals (compare purple bars), but the converse in WU-vaccinated individuals (pink bars).

A characteristic of humoral immunity is that while antibody responses can be boosted by repeated vaccination, the antibody titer saturates when immunity is high and subsequent vaccinations lead to only very modest increases in antibody titers, as is shown in both in the clinical data for CoV-2 and model simulations (Fig 2B) (*8, 30*). We note that in our model, the saturation in the magnitude of the responses occurs due to antibody binding to an epitope sterically preventing B cells specific for that epitope from binding to and being stimulated by that antigen (*24, 25*). This saturation in antibody titers has also been widely observed for other pathogens such as influenza (*24, 31, 36*).

Immune responses get more complex when individuals are exposed to different virus variants or vaccines. These complexities have been discussed in the context of OAS following infections with different strains of influenza. OAS also plays an role for CoV-2 infections, and this is seen in the clinical dataset described by Khan et al. (*29*) (left panel of Fig 2C). Khan et al. show measured antibody titers to both the WU and OM variants in two human cohorts who were infected by the OM (BA1 variant) virus. The first cohort comprised naïve individuals, and the second comprised individuals previously immunized with two doses of WU-vaccines. Vaccinees showed boosting of both WU and OM antibody titers compared with naïve individuals. Interestingly, the WU-vaccine also imprinted responses to the WU-variant, and following OM-infection, these responses reached higher titers compared with antibodies to the OM variant. This is a signature of OAS, and our model reproduces a similar pattern as shown in Fig 2C.

### Model explains the experimental vaccine study of Gagne et al

The most comprehensive and elegant study of boosting by vaccines with new variants are studies which followed vaccination of previously immunized individuals with the original-vaccine versus the updated vaccine (*26*–*28*). We focus on the OM-vaccine study by Gagne et al. (*28*) as the pattern of boosting observed was very similar to the studies based on the Beta variant (*26, 27*).

We used the model to simulate the experiments of Gagne et al., focusing on the responses to the WU and OM viruses (responses to other variants such as Beta and Delta fall in between the responses to WU and OM, as might be expected). Primates were first immunized with two doses of the WU-vaccine and antibody titers were allowed to wane for just under a year. The authors then compared how vaccination #3 with the WU-versus the OM-vaccine boosted responses to both WU and OM virus variants. As mentioned earlier and shown in Fig 3A, Gagne et al. show that the initial two vaccinations (WU-WU) induce higher titers to WU than OM, and that the subsequent vaccination #3 with either WU- or OM-vaccines induce very similar fold-increases in the antibody titers to both WU and OM viruses.

**Fig 3:**
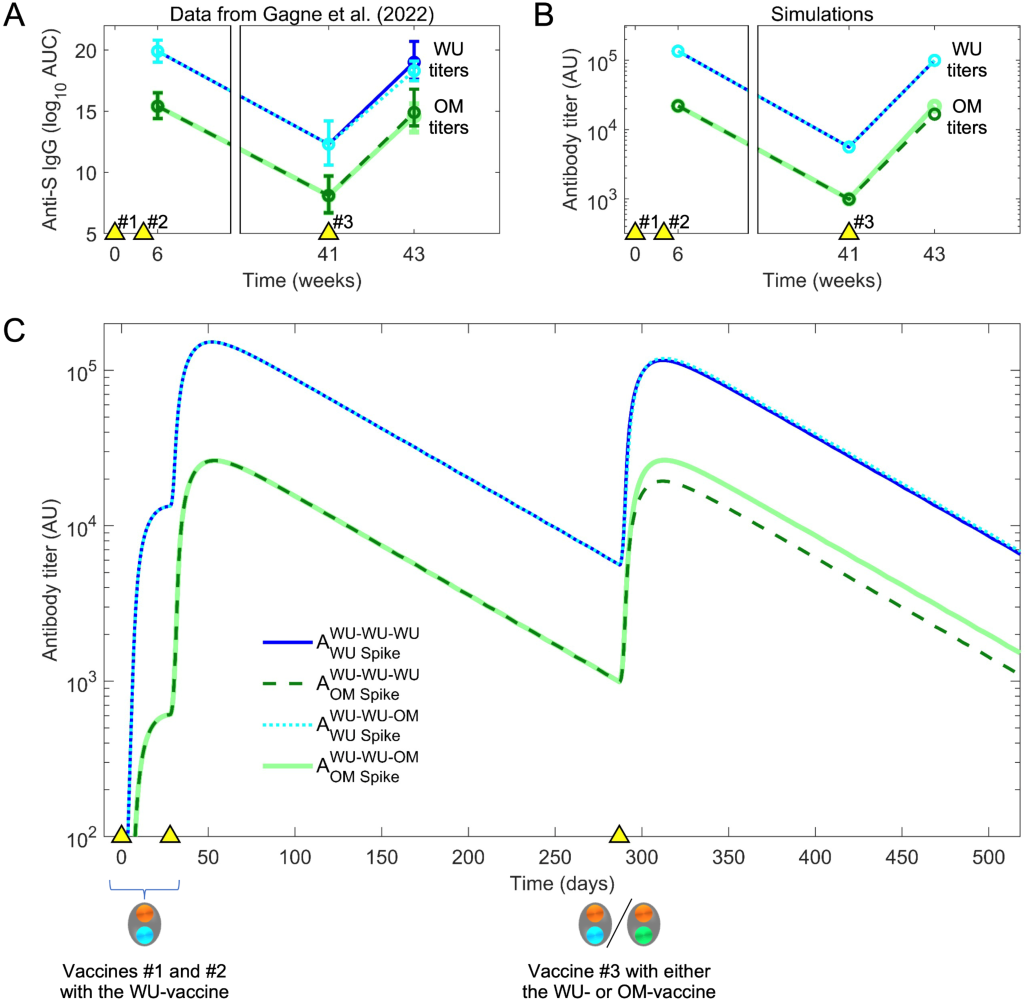
Simulation of the experimental data for boosting. **(A)** Individuals were vaccinated at times indicated by the yellow triangles), initially with the WU-vaccine on days 0 and at 28. The authors found that a second boost (vaccination #3) at day 287 (week 41) with either the WU- or OM- vaccine resulted in comparable titers to the OM virus two weeks later. This trend is captured by the model. **(B)** We show this in more detail by reproducing the results of Gagne et al. (*28*), which shows antibody titers at three timepoints: just after the second vaccination, prior to the third vaccination and 14 days following the third vaccination. The top plot shows that in the experiments of Gagne et al., vaccination #3 with WU- or OM-vaccines caused a similar increase in the titer of antibodies to the OM variant (the color coding is the same as in the legend for panel C). This is consistent with the changes in titers of antibodies to the OM variant in our simulations as indicated in the lower plot **(C)**.

Our model simulations generated the pattern observed experimentally (Fig 3A), and simulations are shown in Fig 3B and C. We then used the model to explore what gives rise to these results. At first glance, there are two surprising observations. First, vaccination #3 with the OM-vaccine does not elicit higher antibody titers to OM than vaccination #3 with the WU-vaccine. Second, vaccination #3 with the OM-vaccine boosts the titer of antibodies to the WU-virus to the same extent as vaccination #3 with the WU-vaccine. From the simulations, we notice that the first observation arises as a consequence of the relationship between the final titer, precursor frequency, and fold boost. Clearly, the final titer equals the product of the precursor frequency and the fold boost. Vaccination #3 with OM (which is the first exposure to OM) results in a significant clonal expansion of B cells unique to OM. However, since the precursor frequency of these cells prior to this immunization is low, the final titer of the response to unique epitopes on OM is relatively modest. In contrast, the precursor frequency of the response to conserved epitopes is high, and even though the fold boost is smaller than that to the epitopes unique to OM (due to epitope masking), these cross-reactive responses form most of the total OM-specific response (see Fig 3B, C).

The model also recapitulated the second observation, namely that vaccination #3 with the OM-vaccine induced similar increases in antibody titers to WU as WU-vaccination #3. This is due to the OM-vaccination stimulating responses to the WU epitopes despite their lower affinity, which is consistent with the explanation of original antigenic sin proposed earlier (*23*).

The model thus shows that though the titer of antibodies to the OM epitope is similar following immunization #3 with either the WU- or OM-vaccines, there are important differences. Vaccination #3 with the OM-vaccine results in a modest increase in OM-specific B cells and antibodies. While these form a small fraction of the total response to OM, we show next that they may have a profound effect following subsequent vaccinations or infections with OM.

### Model predicts scenarios that reveal the benefit of updating the vaccine

We now use our model simulations to examine what would occur if we were to give additional vaccinations (#4 and #5) with OM versus WU. The results are shown in Fig 4A. We see that while the size of the OM response following vaccination #3 is similar whether the OM- or the WU-vaccine is used (the first two bars of the bar plot on the right in Fig 4A), the same does not hold following subsequent vaccinations. Additional OM vaccinations (#4 and #5) result in progressively higher antibody titers to OM compared with a scenario where all vaccinations are with the WU-vaccine. The simulations shown in Fig 4B show that the higher OM-specific response following vaccination #4 & #5 with the OM-vaccine arise due to the generation of antibodies to epitopes unique to OM. These predictions can be experimentally tested if the experimental design of Gagne et al. and similar studies on the Beta variant had included at least one further vaccination (#4). We would expect similar results if exposures #4 and #5 were infections rather than vaccinations. In summary, our model predicts that if vaccination #3 is followed by subsequent vaccinations or infections with the OM variant, it will result in a much higher titer of OM-specific antibodies compared with a scenario where these vaccinations are with the WU-vaccine.

**Fig 4:**
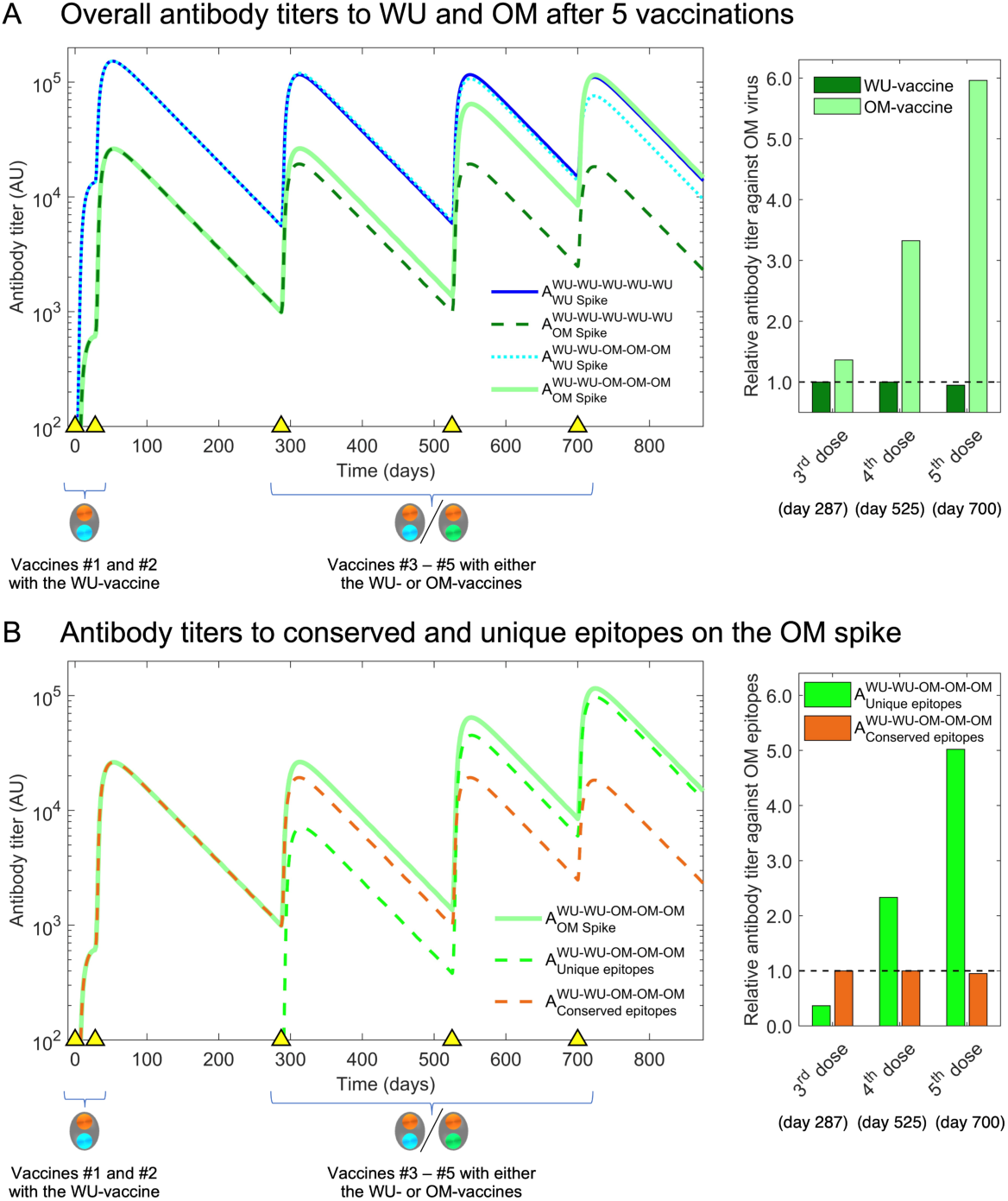
Simulation of third and fourth boosts with the OM-vaccine show generation of higher titers to OM antigen than boosting with the WT-vaccine. **(A)** We plot antibody titers to WU and OM (subscripts) after WU-WU-WU-WU-WU and WU-WU-OM-OM-OM vaccinations. Titers to OM are similar after vaccination #3 with the WU-vaccine and OM-vaccine (solid light green and dashed dark green lines). However, a further vaccination #4 reveals substantially higher antibody titers to OM when the OM-vaccine is used rather than the WU-vaccine (compare solid light green and dashed dark green lines). The bar graph at the right shows that vaccinations #3-#5 are with the OM-vaccine (light green bars) result in much higher antibody titers to OM compared to when the WU-vaccine is used (dark green bars). **(B)** We plot antibody titers to the OM-specific versus conserved epitopes following WU-WU-OM-OM-OM vaccination. We see that the overall increase in titers to OM (light green line) following vaccinations #4 & #5 arises from increases in the OM-specific antibody titer (dashed green line) and responses to the conserved epitopes do not increase (dashed brown line). This is shown in the bar-plot to the right where we see the antibody titer to the OM-specific epitopes (green bars) and shared epitopes (brown bars).

### Predictions are consistent with data for influenza vaccination

The strongest independent support for the prediction that two vaccinations with a new virus strain is needed to reveal the boosting of antibodies to new epitopes comes from influenza H5N1 studies. In Fig 5, we show clinical data from an earlier study (*31*) for immunization with an influenza H5N1 vaccine. Volunteers were immunized with two doses of the hemagglutinin (HA) envelope protein from the H5N1 strain of influenza (which had not circulated in the human population). The HA protein of H5N1 has head and stem domains. The head of H5N1-HA is novel and very different from that of currently circulating influenza strains, while the stem shares conserved epitopes with influenza H1N1, which is circulating in the human population and to which individuals have prior immunity. We see that the first dose of the H5 vaccine results in an increase in the antibody to the shared stem region of HA, and little discernible increase in antibody to the new head region of HA (left panel of Fig 5). However, the situation is reversed following the second immunization with H5. A booster with the H5 vaccine results in substantial increase in the titer to the head of H5, but little further increase in titers to the stem of HA (left panel of Fig 5). This is consistent with the results of our model (see right panel of Fig 5) and supports the hypothesis that generating high antibody titers to novel antigenic sites on a virus protein that exhibits antigenic changes requires two immunizations.

**Fig 5:**
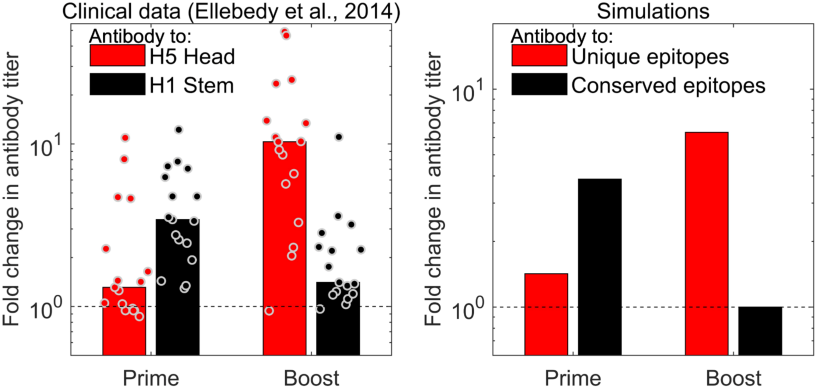
The model captures the observations for influenza vaccination. The left plot shows data for the fold change in antibody titer to epitopes on the head (red) and stem (black) of influenza hemagglutinin (HA) antigen following prime and boost with a H5 vaccine. The first immunization with H5 results in a larger fold increase in antibodies to the conserved stem (shared with H1 viruses), and a significant fold-increase in antibody titers to the unique head epitopes is only seen following the second H5 immunization (boost) (*31*).

## Discussion

Vaccination has played a critical role in the control of the CoV-2 pandemic worldwide (*1, 2*). However, a combination of waning immunity and virus evolution has resulted in large waves caused by new virus variants, in particular the Delta and Omicron variants, that partly evade immunity elicited by the vaccine (*2, 16, 17*). The question then is, when do we need to modify the vaccine to match the circulating virus variant?

Understanding the dynamics of immunity to CoV-2 and influenza are particularly challenging because pre-existing immunity from earlier vaccinations and infections impacts the outcome of subsequent exposures to new virus variants (*20*–*25*). The utility of computational models such as the one we use is their ability to explain complex outcomes that arise from the interactions between multiple factors. The integration of computational modeling to recapitulate patterns observed in multiple datasets can thus play an important role, and ideally should be done in an iterative manner where the models are used to understand the existing data and propose experimental tests that can allow rejection or refinement of the models.

The most important findings of our paper arise from computational modeling of the patterns observed in the elegant experimental study of Gagne et al. (*28*), which compared how the original WU-vaccine versus an updated OM-vaccine boosts immunity to the currently circulating OM virus. Surprisingly, their results showed that WU-WU-WU vaccination was as effective as WU-WU-OM as measured by antibody titers to OM, suggesting that it was not necessary to update the vaccine at the current time. We use mathematical models to address the following: what accounts for this observation, what are the consequences for subsequent immunizations or infections, and how can the model be empirically tested?

Our model suggested that WU-WU-WU and WU-WU-OM result in similar antibody titers to OM because this response is dominated by relatively large secondary (or recall) responses to shared epitopes common to OM and WU. The magnitude of this secondary response obscures the much smaller primary response to new epitopes on OM that occur for the first-time following vaccination #3 with the OM-vaccine (but not with the WU-vaccine).

We then used our models to forecast what would happen if vaccination #3 was followed by further vaccinations or infections. We found that repeated boosts (#3, #4, #5) with OM resulted in much higher titers of antibodies for epitopes unique to OM, and this resulted in a much higher overall titer to OM. Our models thus predict that repeated vaccinations with the updated vaccine are needed to enhance the responses to the new epitopes present in the antigens of new variants. Furthermore, our model suggests a key experiment to allow validation or rejection of the model. The key experiment involves giving one additional vaccine dose (#4) with OM to the primates used by Gagne et al. The model predicts that the group getting WU-WU-OM-OM vaccinations will have much higher antibody titers to OM than the group getting WU-WU-WU-WU. We would expect a similar result (much higher antibody titers to OM) following natural infection with the OM virus after WU-WU-OM vaccination.

Based on this finding, we suggest the general prediction that most of the response generated following the first dose of a vaccine updated to match a new virus variant consists of antibodies specific to the shared antigens, and that high titer responses specific for epitopes unique to the new variant are revealed only following a second immunization with the same vaccine. There may be additional advantages to updating the vaccine to match new virus variants. In particular, it allows the antibody response of individuals to better match future variants that arise from the current OM variant. These variants may correspond to the newly arising lineages of OM (e.g., BA2, BA4, BA5), and antibody responses generated by two doses of the OM-vaccine would be expected to have higher titers to these new variants than if the WU-vaccine were used. Finally, we note that it may be worth considering giving two doses of updated vaccines to vulnerable individuals, not only for CoV-2 but potentially also for influenza.

We now briefly mention several caveats pertaining to our study. At the current stage, we have intentionally used a relatively simple model that focuses on the magnitude of the antibody response following WU- and OM-vaccination. This is because at present, data on the dynamics following immunization and boosting is largely limited to titers of antibodies (*6, 8, 37*–*40*), serum biomarkers (*8, 37, 38*), and the virus inoculum (*41, 42*). We have much more limited data on the dynamics of different populations of cells responsible for the generation of humoral immune responses in the lymph nodes (*39, 43*). These would include different populations of dendritic cells, follicular CD4 T cells, as well as different populations of B cells and plasma cells (*33, 34, 44*–*50*). Further complexities specific to CoV-2 include the spatial aspect of infections of the respiratory tract (*51*–*54*) as well as the dynamics of production and distribution of antigen by mRNA based vaccines (*55*) as well as infections. As more data becomes available, it will be possible to construct more nuanced and refined models of the dynamics of humoral immunity as well as affinity maturation (*56*–*62*). Other directions that could be taken include modeling how protection is lost as the antibody titers elicited by the different immunizations wane. Gagne et al. showed that shortly after vaccination #3, both vaccines elicited similar levels of protection following virus challenge, and it will be important to know if and how this protection declines over time as antibody titers wane (*7, 63, 64*). In particular, we would like to know if the protection against OM infections generated by WU-WU-OM-OM would decline slower than protection following WU-WU-WU-WU. Furthermore we would like to evaluate this for different components of protection, namely, protection from infection versus protection from severe disease (*65*). Another direction is to explore the roles of CD8 T cells (*66*–*68*), particularly those specific to the CoV-2 nucleocapsid protein (*69*) and other viral proteins which may be conserved across CoV-2 strains and might thus play a valuable role in inducing potent cross-reactive immunity.

In summary, the current study uses models to explore some of the complexities associated with choosing when to update the CoV-2 vaccine to match antigenic changes in the virus. Model simulations explain the outcomes of multiple studies of boosting of immunity to CoV-2 and generate qualitatively robust predictions that have implications for determining when to update the CoV-2 vaccine. Based on our results, we suggest that it is not sufficient to monitor the level of immunity to the new variant after a single boost, but that further vaccinations with the updated vaccine should be administered in studies that evaluate the benefit of updating vaccines. This general conclusion may also be relevant for the boosting of immunity to other respiratory viruses such as influenza. An important function of models is that they not only guide the design of vaccination regimes, but also that they are falsifiable, and we have suggested experimental tests that can either confirm or reject the model. Applied to the current debate on updating the CoV-2 vaccine, we propose that a second boost with the OM vaccine be incorporated in studies would result in substantially higher OM-specific antibody titers than if the WU vaccine strain were used.

## Materials and Methods

As mentioned in the text, we extend a multi-epitope model developed earlier (*25, 32*) to consider the dynamics of boosting responses to new strains of influenza. As mentioned in the Results, the model has the following extensions. First, we incorporate two different vaccines, the WU- and the OM-vaccines. Second, we incorporate differences in the boosting of naïve and memory cells to antigenically altered epitopes that underlie the phenomenon of original antigenic sin (*23*). We now describe the model in detail.

Because the available longitudinal data focuses on antibody titers, we use a minimal model that considers 2 vaccine antigens for the WU-vaccine and the OM-vaccine. The antigens WU and OM each have two types of epitopes (Fig 1): the ‘C’ epitopes are conserved across both WU and OM, and the ‘W’ and ‘O’ epitopes are unique to the WU and OM respectively. We let the ratio of conserved to unique antigen epitopes equal *m*: *n* (*m* = 1, *n* = 5; results are qualitatively similar for other values of *m* and *n*). Binding of antibody to the different epitopes on the antigen gives us antigen states as shown in Fig 1. We consider different states for antigen with antibody bound to antigen, for example OM_co_ and OM_xo_ represents OM antigen with no antibody bound (both C and O epitopes free) and OM antigen with antibody bound to the C epitope, respectively. The model also keeps track of B cells B_c_, B_w_ and B_o_ which make antibodies A_c_, A_w_ and A_o_ specific for C, W and O epitope sites, respectively. We use the usual mass action terms for binding of antigen to antibody. B cells are stimulated by cognate antigen (antigen with the relevant epitope free). We further allow previously stimulated (but not naïve) B cells to be stimulated by the altered epitope at a low rate. The latter is a mechanism for original antigenic sin described previously (*23*) and is also validated by the ability of the model to recapitulate CoV-2 boosting data by Khan et al. (2022) shown in Fig 2C. We use standard mass action terms for binding of antibody to antigen and a saturating dose response function for the stimulation of B cells (*25, 32*). The relevant equations for the response to the WU antigen are below (similar equations for the response to the OM antigen are not shown).

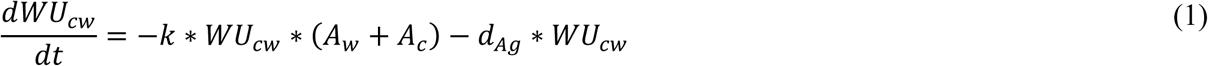

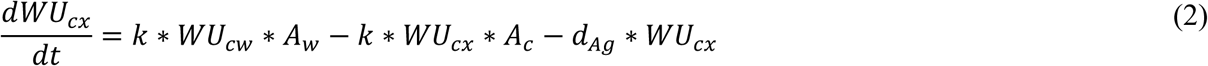

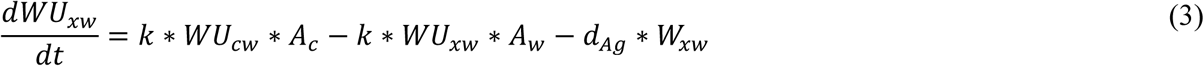

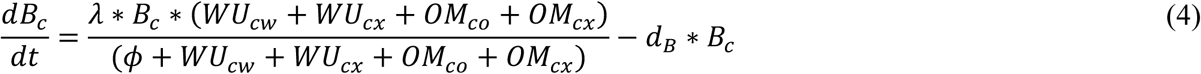

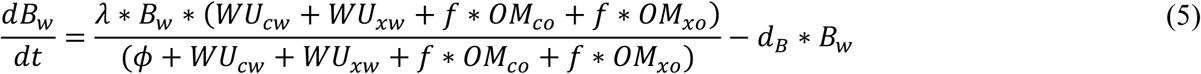

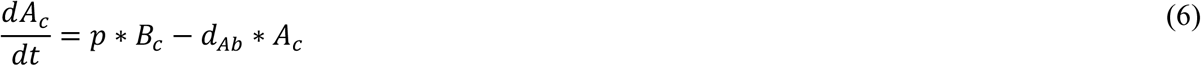

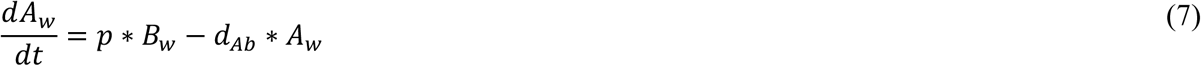

**Table 1:** Parameter values employed in the model. Parameter values are similar to our previous model (*25*). We note that *s* is scaled concentration units, and the initial concentration of B cells is rescaled to 1.

| Model parameter | Symbol | Units | Value(s) |
| --- | --- | --- | --- |
| Rate constant for antibody-antigen binding | $k$ | $s^{-1}day^{-1}$ | 0.0005 |
| Decay rate of free and bound antigen | $d_{Ag}$ | $day^{-1}$ | 1 |
| Decay rate of antibody | $d_{Ab}$ | $day^{-1}$ | 0.1 |
| Maximum proliferation rate of B cells | $\lambda$ | $day^{-1}$ | 1 |
| Antigen for half- maximal proliferation of B cells | $\phi$ | $s$ | 100 |
| Antibody production rate<br>(rescaled so that Antibody=B cell at steady state) | $p$ | $day^{-1}$ | 0.1 |
| Decay rate of B cells | $d_B$ | $day^{-1}$ | $\ln(2)/47$ |
| Fraction stimulation of B cells in secondary responses by non-homotypic antigen | $f$ | — | 0.075 |
| Antigen dose for vaccinations #1 and #2 | — | $s$ | $10^5$ |
| Antigen dose for vaccinations #3, #4 and #5 | — | $s$ | $0.5 \times 10^5$ |
| Time of vaccinations #1 – #5 | — | $day$ | (0, 4, 41, 75, 100) * 7 |
| Ratio of conserved to unique antigen epitopes | $m:n$ | — | 1:5 |

## Abbreviations

WU: Wuhan
OM: Omicron
WU-vaccine: the original mRNA-1273 vaccine encoding the ancestral Wuhan strain (WU virus) spike
OM-vaccine: the updated mRNA-Omicron vaccine encoding the Omicron strain (OM virus) spike
OAS: original antigenic sin

## Acknowledgements

We acknowledge funding from NIH grants U01 AI150747, U01 HL139483, and U01 AI144616.

## Author Contributions

R.D.: Conceptualization, Methodology, Software, Formal analysis, Data curation, Writing – original draft, Writing – review and editing, and Visualization. S.L.L.: Methodology, Formal analysis, Writing – review and editing. C.D.: Methodology, Formal analysis, Writing – review and editing. V.Z.: H.A.: Formal analysis, Writing – review and editing. H.A.: Formal analysis, Writing – review and editing, and Visualization. R.A.: Conceptualization, Methodology, Formal analysis, Writing – original draft, Writing – review and editing, Visualization, Supervision, and Funding acquisition.

## Data and code availability

No new data or original code were reported in this paper. All model simulation and plotting codes and any additional information required to reanalyze the data/simulations reported in this paper are available from the lead contact upon request.

